# Probing solution structure of the pentameric ligand-gated ion channel GLIC by small-angle neutron scattering

**DOI:** 10.1101/2021.04.10.439285

**Authors:** Marie Lycksell, Urška Rovšnik, Cathrine Bergh, Nicolai T Johansen, Anne Martel, Lionel Porcar, Lise Arleth, Rebecca J Howard, Erik Lindahl

**Affiliations:** Department of Biochemistry and Biophysics, Science for Life Laboratory, Stockholm University, Stockholm, Sweden; Department of Applied Physics, Science for Life Laboratory, KTH Royal Institute of Technology, Stockholm, Sweden; Structural Biophysics, X-ray and Neutron Science, The Niels Bohr Institute, University of Copenhagen, Copenhagen, Denmark; Institut Laue-Langevin, Grenoble, France

## Abstract

Pentameric ligand-gated ion channels undergo subtle conformational cycling to control electrochemical signal transduction in many kingdoms of life. Several crystal structures have now been reported in this family, but the functional relevance of such models remains unclear. Here, we used small-angle neutron scattering (SANS) to probe ambient solution-phase properties of the pH-gated bacterial ion channel GLIC under resting and activating conditions. Data collection was optimized by inline paused-flow size-exclusion chromatography, and exchanging into deuterated detergent to hide the micelle contribution. Resting-state GLIC was the best-fit crystal structure to SANS curves, with no evidence for divergent mechanisms. Moreover, enhanced-sampling molecular dynamics simulations enabled differential modeling in resting versus activating conditions, with the latter corresponding to an intermediate ensemble of both the extracellular and transmembrane domains. This work demonstrates state-dependent changes in a pentameric ion channel by SANS, an increasingly accessible method for macromolecular characterization with the coming generation of neutron sources.

## 1 Introduction

Pentameric ligand-gated ion channels (pLGICs) mediate electrochemical signal transduction in most kingdoms of life. In metazoa, these receptors populate postsynaptic membranes, where they open upon binding neurotransmitters to control the firing of action potentials; accordingly, human pLGICs are targets of anesthetic, anxiolytic, and other important classes of drugs [1]. Several prokaryotic pLGICs have also been identified, possibly involved in chemotaxis or quorum sensing [2]. Despite their physiological significance, our picture of the pLGIC gating landscape is still incomplete, and we only have limited knowledge e.g. about the structural dynamics of conformations at room temperature or intermediate states, both of which are essential for understanding of fundamental channel biophysics as well as potential drug development.

The proton-gated cation channel GLIC, from the prokaryote *Gloeobacter violaceus*, has been a popular model system for structure-function studies in this superfamily [3]; it accounts for 40% of all pLGIC structures in the Protein Data Bank. This channel is pH-gated, and its structure has been solved at both high and low pH in apparent closed and open states, respectively [4, 5, 6, 7]. Additional states have been determined in the presence of engineered mutations or lipids, possibly representing gating intermediates [8, 9, 10, 11, 12]. Each channel is comprised of five identical subunits, each of which contains an extracellular domain (ECD) consisting of ten *β*-strands, and a transmembrane domain (TMD) of four *α*-helices, **Figure 1A**. Relative to the closed structure, apparent open structures of GLIC undergo contraction in the ECD towards the central conduction pathway, as well as outward tilting of the pore-lining helices [6], **Figure 1B**. Recent structures of eukaryotic homologs including GluCl, GlyR, GABA_*A*_R, and 5-HT_3_A receptors indicate similar gating transitions, although the specific geometries and amino-acid contacts are slightly different [3, 13].

**Figure 1:**
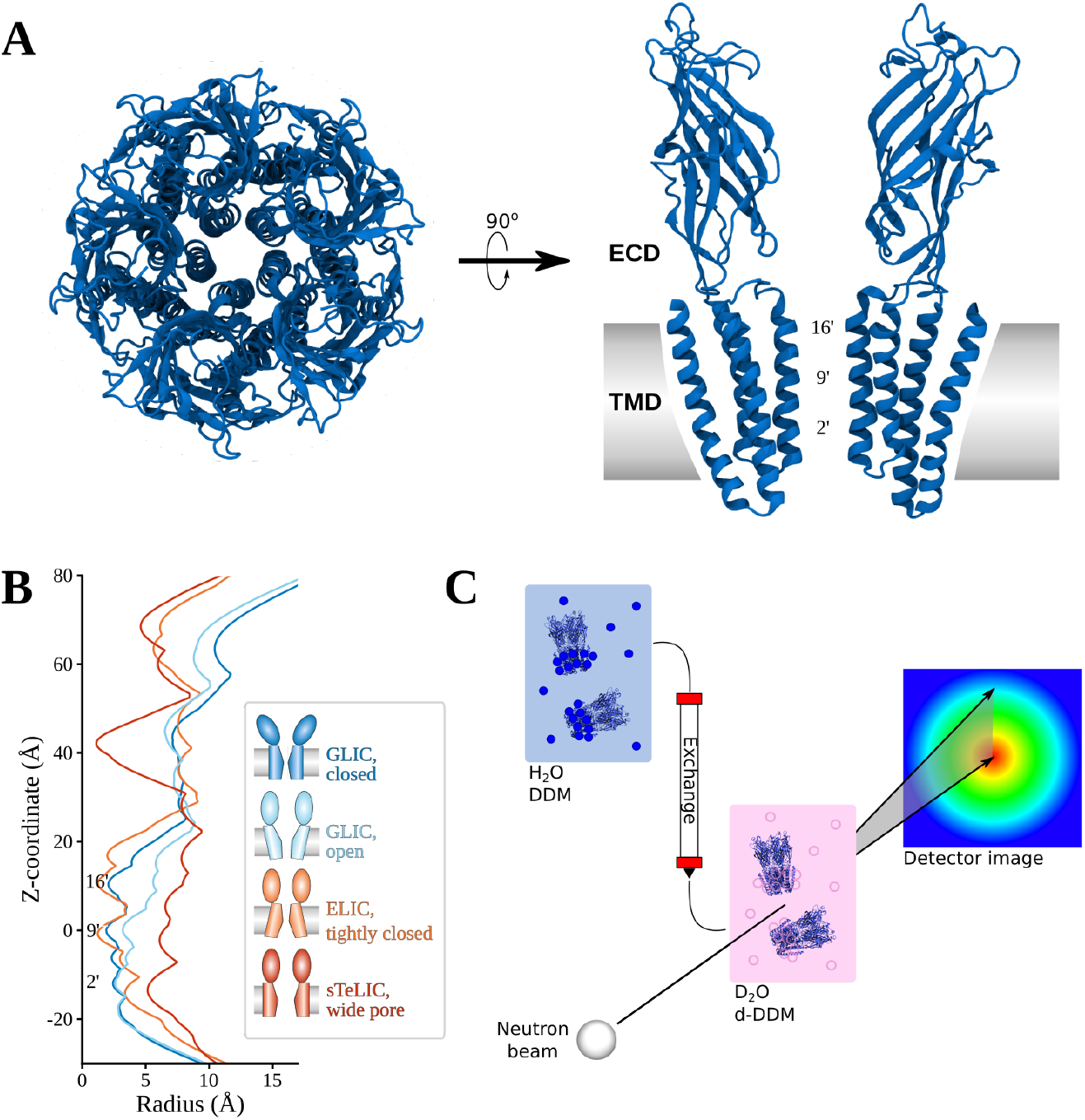
Proposed structures and experimental approach for characterizing solution-phase GLIC by SANS. **A)** Extracellular view of GLIC (PDB ID: 4NPQ, closed), and two subunits seen from the membrane. The ECD in each subunit contains ten β-strands, and the TMD four helices. Constriction points are labeled with the common prime notation for the pore lining helix of pLGICs. **B)** Pore profiles calculated using Hole [14] for closed GLIC (PDB 4NPQ, dark blue), open GLIC (PDB 4HFI, light blue), tightly closed ELIC (PDB 3RQU, orange) and wide open sTeLIC (PDB 6FL9, red). Relative to the closed state, open GLIC has a wider pore at 9’ and 16’, while the top of the ECD has moved inwards. ELIC and sTeLIC are similar in the ECD where both have a lower radius than GLIC, while the former is even more collapsed than closed GLIC in the upper part of the TMD, open sTeLIC has a wider pore than the other structures. **C)** Schematic of the SEC-SANS experiment: A sample in H_2_O-based buffer with DDM detergent is loaded onto a size exclusion column running a D_2_O-based buffer with match-out deuterated DDM (d-DDM), resulting in the protein being exchanged to the D_2_O and d-DDM environment where the detergent is indistinguishable from the buffer in terms of neutron scattering. The SANS cell is in-line with the size exclusion chromatography, leading to measurements directly following size-based separation of the sample, which avoids sample aggregation.

Although X-ray structures of GLIC offer models both of stable and intermediate conformations, their correspondence to specific functional states on the gating pathway is still a matter of discussion. Structures of other bacterial family members suggest there are also alternative conformations in the pLGIC landscape, including a more tightly closed form of the pore in numerous structures of the plant-pathogen channel ELIC [15, 16, 17], and a dramatically expanded pore in the symbiote channel sTeLIC [18], **Figure 1B**. Both the ELIC and sTeLIC structures are contracted in the ECD compared to either closed or open GLIC, indicating further conformational diversity in this domain too. This could suggest some channels (e.g. GLIC) might be able to exist in conformations not yet observed experimentally, or that features such as crystal packing, experimental conditions or the liquid nitrogen freezing might have biased the conformational distribution. Indeed, the vast majority of GLIC structures to date contain an activated ECD conformation; the few resting-state structures available were reported to notably lower resolution [6, 7]. Biophysical techniques including electron paramagnetic resonance spectroscopy [19, 11] and atomic force microscopy [20] have suggested GLIC can sample additional states outside the crystalline context, emphasizing the importance of alternative structural approaches to model pLGIC conformational cycling in solution.

The solution structure of a protein can be probed using small-angle scattering, in which a sample’s distinctive scattering of X-rays or neutrons allows modeling of its population-average structure, **Figure 1C**. Small-angle neutron scattering (SANS) was applied as early as 1979 to the muscle-type nicotinic acetylcholine receptor, a pLGIC extracted from native tissue of electric rays, yielding low-resolution predictions of molecular volume, radius of gyration, and overall size and shape [21]. Thanks to the differential scattering of neutrons by isotopes of hydrogen, it is possible to hide the micellar environment of a membrane protein by matching the contrast of the detergent and the solvent. This has lead to the development of *match-out* deuterated detergent [22] and nano-discs [23], where the contrast is matched to pure heavy water (D_2_O). This strategy has recently been applied to probe AMPA-receptor, ABC-transporter, ATPase, photosystem-I, and translocon structures in solution [22, 24, 25, 26]. However, given the relatively low resolution of SANS the method has traditionally not been able to resolve minor structural changes between states due e.g. to gating in ion channels in general and pLGICs in particular.

Another challenge is that membrane proteins can be highly prone to aggregation, which will lead to major distortion in the SANS spectrum, but this can be minimized with new methods that pass the sample directly to the neutron beam from a size-exclusion chromatography column (SEC-SANS) [27, 28], **Figure 1C**. This approach can be run continuously from column to measuring cell, or with pausing of the flow when the protein peak reaches the cell, enabling measurement at multiple detector distances in a single run. The continuous flow approach has been demonstrated [22, 28, 29], but there is little precedence in the literature for the paused-flow approach.

Here, we took advantage of SEC-SANS in match-out deuterated detergent to characterize solution structures of GLIC, which enabled us to measure scattering curves that exhibit slight differences between resting vs. activating conditions. By calculating theoretical scattering curves, we show that the closed-state X-ray conformation is the single experimental structure that best matches the experimental data, which excludes several alternative states. To better model the scattering data, we further applied enhanced-sampling molecular dynamics (MD) simulations to identify different ensembles of best-fit models in each condition, finding that an intermediate conformation or mixture of states best represents GLIC under activating conditions. This work demonstrates how SANS is able to resolve even subtle conformational changes between functional states of an ion channel at room temperature, the utility of SEC-SANS, and how the integration with MD-based enhanced sampling of flexibility in multiple states enables better matching of the activating conditions SANS measurements than any single experimental structure.

## 2 Results

### Subtle effects of agonist (pH) on GLIC scattering

To investigate the solution structure of GLIC under resting and activating conditions, we first collected paused-flow SANS measurements of purified GLIC at pH 7.5 (resting conditions) and 3.0 (activating conditions), respectively. Guinier analysis to determine the radius of gyration revealed similar molecular radius (38.4 ± 0.2 Å at pH 7.5, and 38.3± 0.2 Å at pH 3.0) and weight estimates (194 kDa for both, with an expected error of 10% from the uncertainty in the protein concentration determination), **Figure 2A**. These values are in line with what is expected from the crystal structures (R_g_ of 37.9 Å for the closed structure, and 37.4 Å for the open), and from the amino acid sequence of the construct (182.7 kDa). For conformations this similar, it is not realistic to discriminate states directly from the overall radius of gyration. However, the scattering curves exhibited slight differences at Q-values around 0.10 to 0.12 Å^-1^ – the region where differences are expected based on theoretical scattering curves calculated from crystal structures. A summary of the structural parameters calculated from the experimental curves is available in **Supplementary Table S1**.

**Figure 2:**
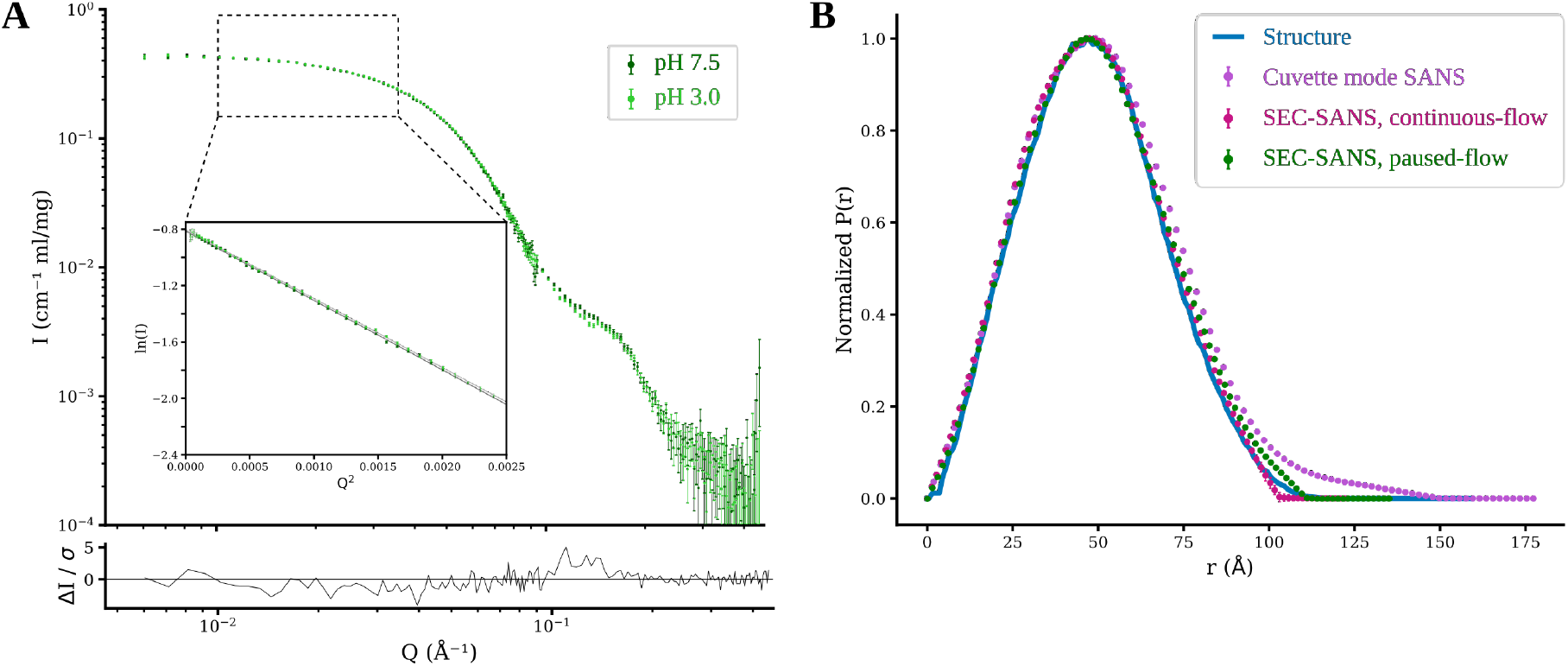
Subtle effects of agonist (pH) revealed by paused-flow SEC-SANS. **A)** SANS experimental data for GLIC measured using paused-flow SEC-SANS at resting (pH 7.5) and activating (pH 3.0) conditions, and the residual between the measured data sets. The overall shape of the experimental data is similar, but a difference is observed in the region *Q* ∈ [0.10, 0.12] Å^-1^. The Guinier plot of the low Q-region is shown in the inset, with the line representing the Guinier fit. **B)** Pairwise distance distributions calculated from the closed GLIC model (blue), and derived from measurements at pH 7.5 using cuvette SANS (purple), continuous-flow SEC-SANS (standard SEC-SANS, pink), and paused-flow SEC-SANS (green). Both continuous-flow and paused-flow SEC-SANS avoid aggregation, unlike the cuvette measurement.

Concerning GLIC measured at pH 7.5, the paused-flow SEC-SANS arrangement - which enabled measurement of the same fraction at two detector distances - was compared with the classical cuvette-SANS configuration, and the continuous-flow SEC-SANS arrangement. Coupling inline size exclusion immediately prior to scattering measurements (SEC-SANS) eliminated the accumulation of low-order aggregates evident in the cuvette measurement. This is obvious in the pairwise distance distribution where there was close agreement between both SEC-SANS data-sets and the distribution expected from the closed atomic model, while the cuvette measurement displayed a low amount of interatomic distances up to 150 Å, **Figure 2B**.

### Solution scattering corresponds to resting X-ray structures

To assess the how the shape of solution-phase GLIC matches putative 3D structures, we first compared our SANS data to scattering curves calculated from atomic models derived from previously reported X-ray structures: GLIC crystallized at pH 7.0 (closed, PDB ID 4NPQ) and 4.0 (open, PDB ID 4HFI), ELIC in the absence of agonist (tightly closed pore, PDB ID 3RQU), and sTeLIC in the presence of agonist (wide pore, PDB ID 6FL9). For the latter two comparisons, to ensure that model fits represented differences in large-scale conformation rather than local sequence differences between channels, we constructed homology models of GLIC based on template structures of ELIC and sTeLIC, respectively. Fitting of these theoretical spectra to the experimental measurements showed that the closed-state GLIC structure exhibited the best fit to the scattering curves under both resting and activating conditions (χ ^2^ goodness of fit of 2.8 at pH 7.5 and 2.6 at pH 3.0), **Figure 3**. The second-best match was obtained with the open structure of GLIC, but this only resulted in moderately good fits, with a χ ^2^ of 7.3 under activating conditions and slightly higher (8.8) for the resting-condition measurement. The models based on the ELIC/sTeLIC homologs resulted in clearly worse fits than either GLIC structure, indicating that the room-temperature solution-phase structure of GLIC indeed is better described by the GLIC conformations observed by X-ray crystallography than by more open or closed conformations observed in related channels. The wide pore and contracted extracellular domain conformation of sTeLIC was a particularly poor model for solution-phase GLIC (χ ^2^ of 22 at pH 7.5 and 20 at pH 3.0).

**Figure 3:**
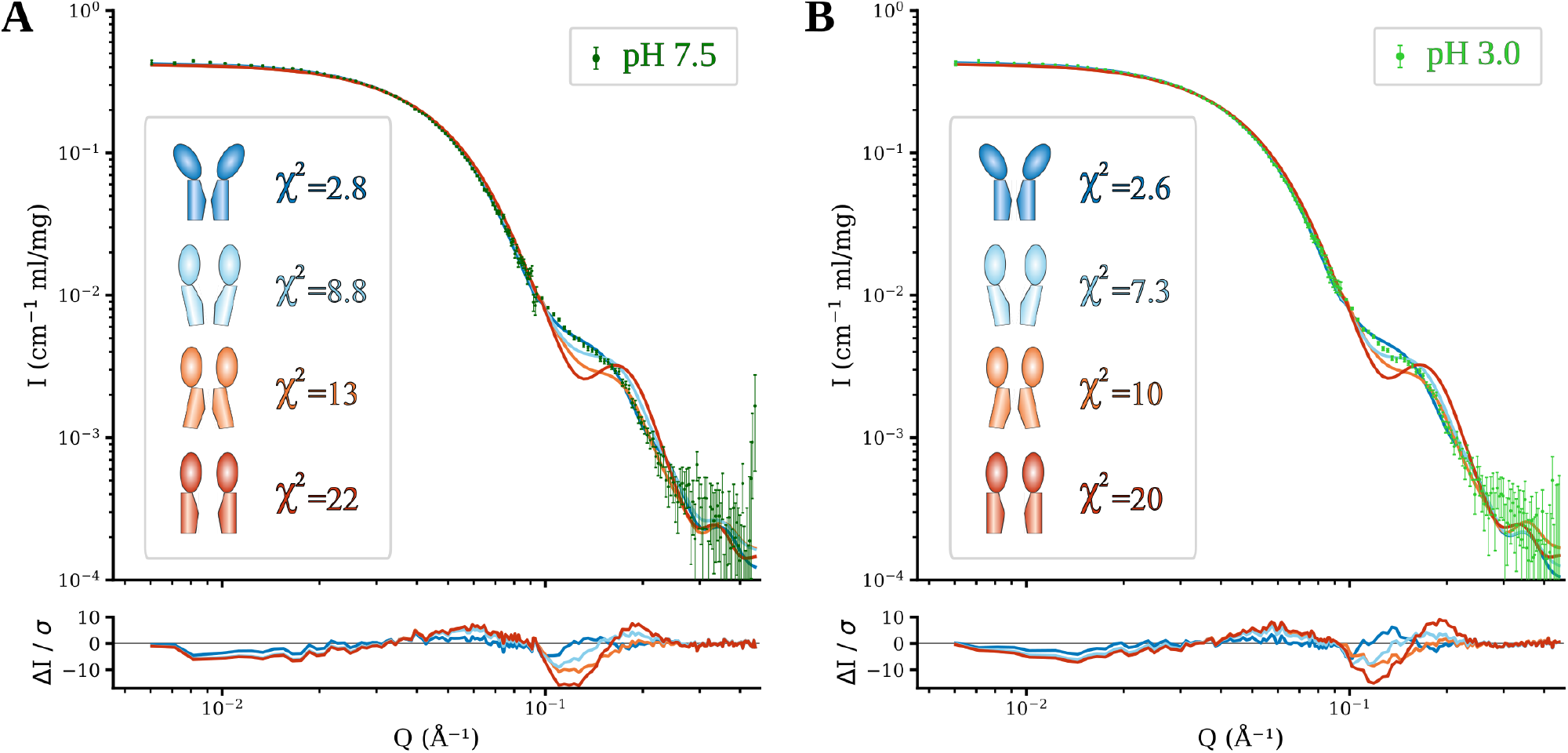
GLIC solution scattering corresponds to resting X-ray structures. Fits of model spectra calculated from crystal structures to SANS data collected at **A)** pH 7.5 (resting conditions) and **B)** pH 3.0 (activating conditions). Based on the goodness of fit (χ^2^), the individual structure that provided the best fit to both measurements was closed-state GLIC, with the second-best (though substantially worse) match between open-state GLIC and activating condition measurements. Any linear combination of theoretical curves made the fit to resting state data worse, while up to 18% of the open structure leads to at least equally good fits to the activating conditions SANS curves, Supplementary Figure S1.

Fitting of a linear combination of the theoretical curves from the closed and open models of GLIC revealed that including any fraction of the open model worsened the goodness of fit to the resting state data, while including more than 18% from the open structure worsened the fit at activating conditions, **Supplementary Figure S1**.

### Molecular dynamics simulations enable modeling of pH effects

Since comparisons to individual X-ray structures did not conclusively discriminate differences between resting and activating conditions in solution, we tested if the data could be better described with sampling of more diverse regions of the conformational landscape. This can in principle be achieved with molecular dynamics simulations; provided the simulations are started in many different regions of the conformational space they should provide an ensemble of conformations that approximate the flexibility and motions present during gating. To generate such an an ensemble, we used coarse-grained elastic network extrapolation between the open and closed structures of GLIC, followed by re-introduction of atoms absent in the coarse-grained extrapolation (side chain reconstruction) and molecular dynamics simulations from these seeds. After side chain reconstruction, energy minimization, and equilibration, the fits showed major improvements over experimental X-ray structures to a best χ ^2^ of 1.4 to the pH 7.5 data and 1.5 to the pH 3.0 data. This was improved just-so-slightly further after 200 ns of production simulation (best χ ^2^ of 1.3 to the pH 7.5 data and 1.4 to the pH 3.0 data), **Supplementary Figure S2**. The best fitting models had theoretical curves in close agreement with the scattering curve, including in the 0.10-0.12 Å^-1^ region, **Figure 4A** and **B**. A summary of the modeling using crystal structures and molecular dynamics simulations is available in **Supplementary Table S2**.

**Figure 4:**
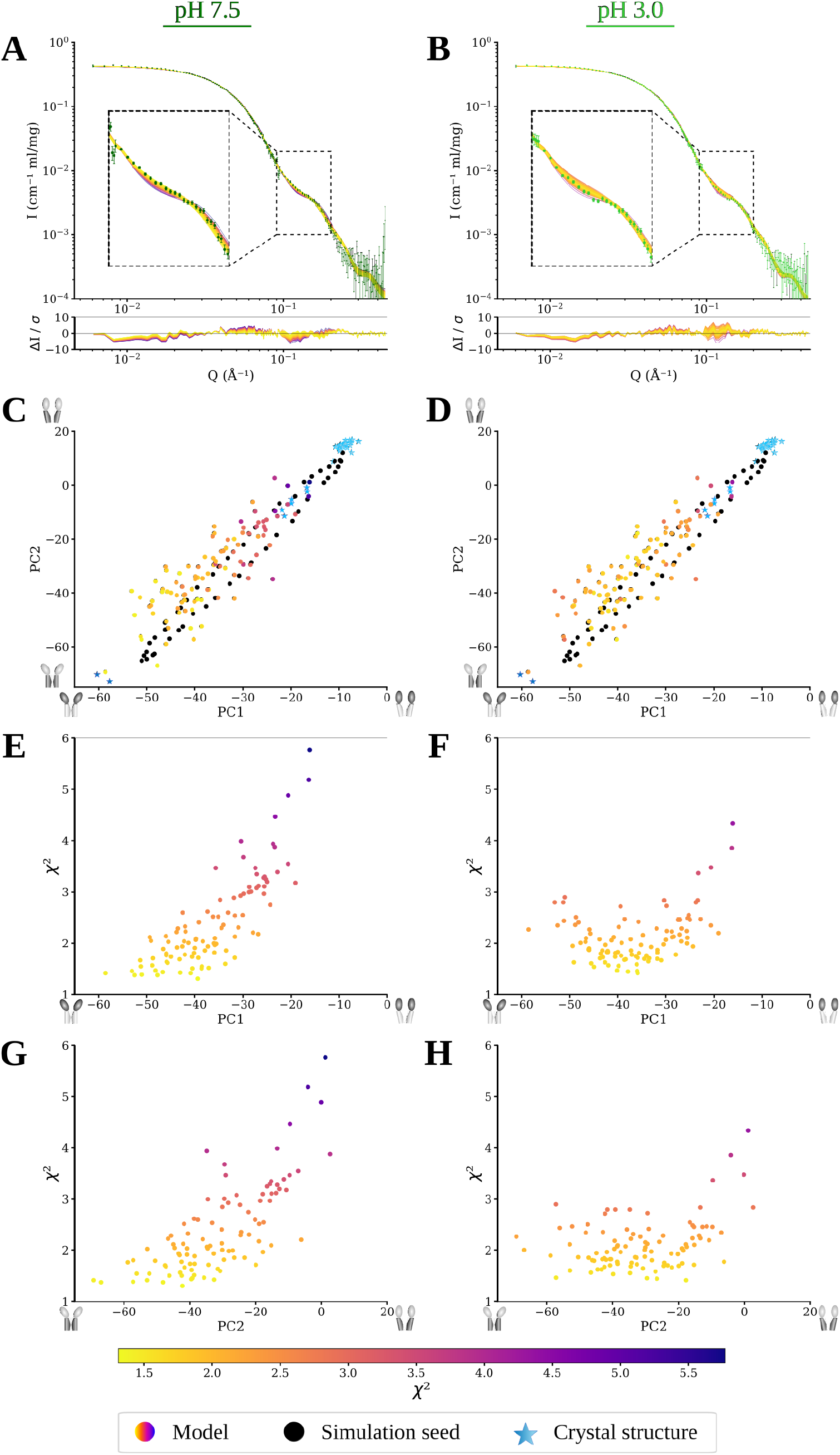
Comparison of molecular simulation ensembles to experimental SANS data at pH 7.5 (left column) and 3.0 (right column). **A)** and **B)** Experimental SANS curves overlaid with theoretical scattering curves from models after 200 ns of molecular dynamics simulations. Colors represent goodness of fit to the experimental curve. The magnified inset corresponds to *Q* ∈ [0.1, 0.2] Å^-1^. **C)** and **D)** Conformations from simulation ensembles projected onto the two lowest principal components, PC1 and PC2, of the landscape constructed from 46 GLIC crystal structures (blue stars), with the main motion captured by each PC illustrated by the dark half of the cartoons on the axes. The energy minimized interpolated conformations used as simulation seeds are shown as black dots. Colored data points represent conformations after 200 ns of molecular dynamics simulations, with shade representing their χ ^2^ goodness of fit to the pH 7.5 (C) and 3.0 (D) experimental SANS data. Panels **E)** and **F)** project all simulation conformations on PC1 along the x-axis, while the y-axis shows χ ^2^ for data at pH 7.5 (E) and pH 3.0 (F). Finally, panels **E)** and **F)** display the same data but instead projected onto PC2 at pH 7.5 (G) and pH 3.0 (H).

Principal component analysis (PCA) can be used to identify which motions in a system explain most of the observed variance. Using the first two principal components defined by 46 experimental GLIC structures [30], models that best fit SANS data under resting conditions largely correspond to the lowest coordinates of both PCs, near the closed X-ray structures, **Figure 4C**. In contrast, for the SANS data from activating conditions the region with the best fitting models was shifted towards the open structure, **Figure 4D**.

The first principal component largely corresponded to the expansion of the ECD, while the second mainly captured the contraction of the upper part of the pore (**Supplementary Video 1 and 2**). By correlating the goodness of fit to each PC, it is possible to assess how these conformational features impact the fit. For the resting state SANS data, the goodness of fit is clearly best (lowest) at the coordinates corresponding to the closed GLIC structure, and it increases almost linearly as the system undergoes the transformation towards the open state both for PC1 (**Figure 4E**) and PC2 (**Figure 4G**). In contrast, at activating conditions the best fit is achieved for ensemble simulation states from roughly halfway between closed and open GLIC states (**Figure 4F,H**). Based on the motions the PCs correspond to, this indicates that under resting conditions the best fit is achieved with a fully expanded extracellular domain and a contracted pore, while an intermediate expansion of the extracellular domain yields the best fit to SANS data from activating conditions, with tolerance for a range of pore conformations, **Figure 5**.

**Figure 5:**
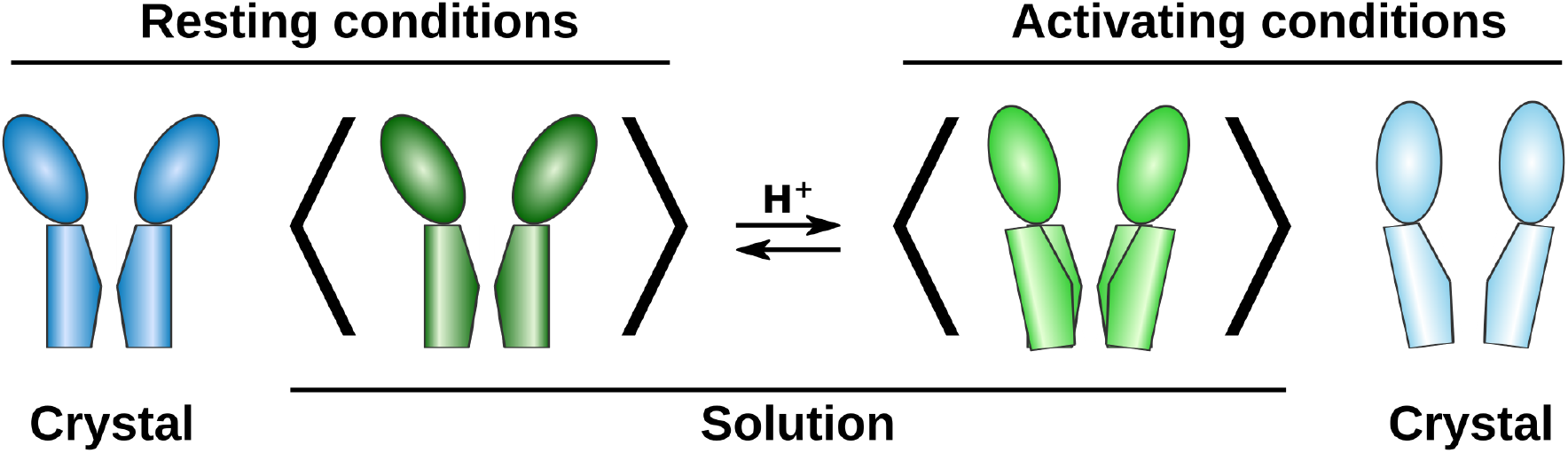
Conceptual models of GLIC conformational distributions. Under resting conditions the average solution structure of GLIC is similar to the closed X-ray structure, while activating conditions appear to favour an intermediate ECD-expansion and a range of possible pore conformations, which could also correspond to a mix of multiple discrete conformations.

## 3 Discussion

By comparative fitting of SANS curves to crystal structures and enhanced-sampling MD ensembles, we have demonstrated that it is possible to use SANS to detect quite subtle conformational changes in pentameric ligand-gated ion channels in solution phase at room temperature, and test structural hypotheses. Using crystal structures of GLIC and related pLGICs, we were able to rule out substantial contributions from putative alternative states containing a contracted ECD and either a wide or tightly closed pore, and we identify the closed GLIC X-ray conformation as the best single-structure representative (χ ^2^ ≤ 2.8) of the average solution ensemble at resting conditions, while the data at activating conditions is compatible with a mix of up to 18% contribution from the open structure. By further sampling conformations along the predicted gating transition and using an ensemble of MD simulation structures, we were able to achieve significantly better fits to both resting and activating condition SANS data, and can clearly discriminate between structural models to fit the data. The differences are mainly evident in the shift of the scattering curves at high Q (>0.1 Å^-1^). This suggests a model for the conformational distribution where the average conformation of GLIC under resting conditions has an expanded ECD and contracted pore, and is well described by the closed X-ray structure (**Figure 5**). Upon activation (low pH), the average conformation of the ECD partly contracts towards the open X-ray structure, while the average TMD conformation shows a broader distribution that could also indicate a mix of multiple discrete states.

Given the relatively subtle differences in GLIC scattering under different pH conditions, the observations in the present study would not have been possible without several recent experimental advances. As recently demonstrated for other membrane-protein families [22, 24, 26], use of deuterated detergent produced scattering data representing the protein alone, simplifying data analysis by allowing the exclusion of micelle parameters. In cuvette mode, GLIC showed evidence of aggregation even when exchanged by gel filtration on the same day, **Figure 2B**. Switching to the SEC-SANS configuration proved to be instrumental in reducing the contribution of apparent aggregates. A drawback of this mode is that continuous SEC-SANS requires the sample to be split over multiple runs in order to measure at multiple detector distances: for GLIC as for many macromolecules, collection at two distances was necessary to obtain both low-resolution information related to sample mass 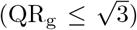 and higher-resolution information related to functional state (Q>0.1 Å^-1^), **Figure 2A**. Despite requiring some technical finesse, manual suspension of SEC flow and temporary relocation of the SANS detector during peak elution enabled measurement with minimal aggregates over an extended Q-range for a single sample load, **Figure 2B**; this “paused-flow” configuration will likely be applicable to other biological macromolecules. With the latest update to the D22 instrument at the Institut Laue-Langevin neutron source, the upgrade to two detector banks will allow measuring the a Q-range from 0.003 Å^-1^ to 0.7 Å^-1^ in a single run without manipulating the detector position during the run. Finally, to sample beyond the available crystal structures, coarse-grained interpolation followed by all-atom MD simulations proved highly efficient in modeling GLIC SANS curves, **Figure 4**. Given the number of interpolation seeds used, even the modest simulations implemented here (200 ns each) could constitute a prohibitive computational cost (20 ms total simulation time) for similar future work. However, we noted a similar quality of fits for relaxed simulation seeds as for simulation endpoints, **Supplementary Figure S2**, which suggests the production runs could possibly be much shorter in many cases.

SANS primarily differentiates structural models on the basis of large-scale conformational changes that affect e.g. the peripheral radius. In GLIC, such transitions primarily involve the contraction or expansion of the ECD. Accordingly, the best-fitting crystal structure at both pH conditions was closed (resting state) GLIC, due to its more expanded ECD, **Figure 3**. Indeed, in linear combinations only a minority of the open GLIC structure was tolerated, **Supplementary Figure S1**. Notably, lattice interactions in pLGIC crystals primarily involve the ECD periphery, as other regions are embedded in or proximal to the detergent micelle; it is possible that more contracted forms of the ECD might be preferentially stabilized by crystal contacts, while a slightly more expanded state predominates in solution. Still, the best-fit MD-simulation models under resting conditions corresponded to the closed crystal structure not only along PC1, which primarily represented ECD blooming; but also along PC2, representing upper-TMD contraction, **Figure 4**. We found no evidence for states comparable to ELIC or sTeLIC in GLIC solution structures. Aside from a contracted ECD, ELIC- and sTeLIC-based models contained tightly closed or wide-open pores respectively, **Figure 1**, which did not appear to contribute based on the distribution of best-fit MD models along PC2; instead, resting conditions corresponded closely to the closed crystal structure, while activating conditions were best modeled by simulation ensemble members roughly halfway between open and closed X-ray structures. The locally closed state of GLIC [8] projects to intermediate values in the principal component analysis, **Figure 4**, a bit closer to the open crystal structure than the optimal region for fit to the activating condition SANS data. The locally closed structures have a contracted ECD like the open structure and a closed pore, but the average solution conformation at activating conditions appears to be somewhat closer to the closed crystal structure in both the ECD and TMD. Thus, our results emphasize expansion of the ECD relative to open GLIC crystal structures, but also highlight moderate pH-dependent changes in the TMD.

Fits to scattering curves under resting conditions (pH 7.5) contrasted with those under activating conditions (pH 3.0). For the latter, SANS was best described by conformations interpolated between resting and open crystal structures. Because SANS measures an ensemble, an intermediate model may represent a distinct dominant state, or the average of a heterogeneous mixture. Indeed, several structural studies have supported the presence of multiple states in equilibrium at low pH. One GLIC variant was crystallized at pH 4 in two partially occupied forms, both with a contracted ECD, but with either a closed or open pore [6]. Atomic force microscopy images have also revealed a mixture at low pH, including states that predominated under neutral resting conditions [20]. We have also recently used cryo-electron microscopy of GLIC to reconstruct multiple particle classes at pH 3, including a dominant state with an expanded ECD and closed pore, and a minority class intermediate between resting and open crystal structures [7]. Due in part to its low conductance in single-channel recordings [31], the open probability of GLIC has not been well established. However, there is precedence in this family for flickering between states even at high agonist concentrations [32, 33] and for open probabilities well below 100% at activating conditions [34, 35, 36, 37]. In recent computational work, we estimated the low-pH open probability of GLIC to 17% using Markov state modeling [38] – notably similar to the tolerated contribution (18%) of the open structure in our linear-combination fits, **Supplementary Figure S1**. There are thus several indications the SANS curve under activating conditions might be due to an average of mixed states, mostly closed, but some similar to open crystal structures. Alternatively, pH-3 conditions could promote a predominant desensitized state, distinct from both resting and open crystal structures. It has been proposed that desensitization mimics the lipid-modulated crystal structure of GLIC, with a contracted ECD and partially expanded pore [11]. Although this would be slightly different from the intermediate ECD-contraction of the average pH-3 structure, contribution from this or another desensitized form might add to the complexity of the GLIC conformational landscape at low pH.

In conclusion, paused-flow SEC-SANS proved effective in determining distinct room temperature solution structures of a key model pentameric ligand-gated ion channel under resting versus activating conditions. The resolution of this approach is still limited by the signal-to-noise ratio at high Q of the SANS experiment. However, this is expected improve by an order-of-magnitude as next-generation neutron setups (e.g. the European Spallation Source ESS) are taken into production the coming years, which will enable lower sample volumes and correspondingly higher concentrations. This should make it a very powerful method for discerning even small conformational transitions of biological macromolecules under realistic solution-phase and room temperature conditions.

## 4 Methods

### 4.1 Protein expression and purification

GLIC was expressed and purified as previously described [5, 39]. In short, a fusion protein of GLIC and maltose binding protein (MBP) in a pET-20b derived vector was expressed in C43(DE3) *Echerichia coli* cells. Cells were inoculated 1:100 into 2xYT media with 100 mg/mL ampicillin, grown at 37° C to a OD_600_ of 0.7, at which time they were induced with 100 mM isopropyl-b-D-1-thiogalactopyranoside (IPTG), and shaken overnight at 20° C. Cells were harvested through centrifugation and lysed by sonication in buffer A (300 mM NaCl, 20 mM Tris-HCl pH 7.4) supplemented with 1 mg/mL lysozyme, 20 μg/mL DNase I, 5 mM MgCl_2_, and protease inhibitors. Membranes were harvested by ultracentrifugation and solubilized in 2% n-Dodecyl-β-D-Maltoside (DDM), following which the GLIC-MBP fusion protein was isolated through amylose affinity purification and size exclusion chromatography in buffer B (buffer A with 0.02% DDM). The fused MBP was cleaved from GLIC using thrombin, and an additional size exclusion chromatography yielded pure GLIC in a buffer B.

### 4.2 Small angle neutron scattering

SANS experiments [40, 41] were performed at the D22 beamline of Institute Laue–Langevin, using three experimental set-ups: cuvette-mode SANS, continuous-flow SEC-SANS, and paused-flow SEC-SANS, see **Supplementary Tables S3 and S4** for detailed sample and collection parameters. Cuvette-mode allowed the full amount of sample to be measured for both detector distances (2m/2.8m, and 11.2m/11.2m) required to achieve the full Q-range, it did not however ensure sample monodispersity. To measure monodisperse samples SEC-SANS [27, 28] was used, running a gel filtration with a Superdex 200 Increase 10/300 column in-line with the SANS measurement. In the continuous-flow measurement the sample was divided over two runs, one for each detector setting (2m/2.8m, and 11.2m/11.2m), and the flow speed decreased to 0.05 ml/min upon peak detection by UV-vis absorbance at 280 nm. In the paused-flow measurements the full sample amount was loaded for a single run, the flow was slowed down to 0.01 ml/min upon peak detection and paused at the peak max to enable the second detector distance measurement (2.8m/2.8m during the run, and 8m/8m while paused). The gel filtration step was used to exchange the sample to D_2_O buffers with match-out deuterated DDM (d-DDM) [22]. For the cuvette-mode experiment this was performed off-line to the SANS experiment and the exchanged sample loaded into 2 mm Hellma quartz cuvettes. All three experimental set-ups were used to measure GLIC at pH 7.5 (150 mM NaCl, 20 mM Tris·HCl pH 7.5, 0.5 mM d-DDM), GLIC at pH 3.0 (150 mM NaCl, 20 mM citrate·HCl pH 3, 0.5 mM d-DDM) was measured using the paused-flow set-up.

Data reduction and buffer subtraction were performed using Grasp version 9.04 [42], utilizing either detector frames preceding the protein peak (standard SEC-SANS) or dedicated buffer measurements for the buffer background (paused-flow SEC-SANS and cuvette SANS). Data were corrected for the empty cuvette and background, and scaled by their transmission and thickness. They were scaled to absolute intensity by direct flux measurement. Concentration normalization of the paused-flow SEC-SANS runs were performed by dividing the scattering intensity with the concentration calculated using the absorbance at 280 nm from the co-recorded chromatograms and the extinction coefficient calculated from the amino acid sequence using ProtParam [43]. In merging data from the two detector distances, data from the detector distance covering low Q (11.2m for continuous-flow and cuvette-mode, 8m for paused-flow) were used up to 0.070 Å^-1^ continuous-flow and cuvette-mode, and up to 0.095 Å^-1^ for paused-flow. Above this, data from 2m (continuous-flow and cuvette-mode) and 2.8m (paused-flow) were used. A small additional constant was subtracted as a final adjustment to the background.

### 4.3 Model preparation

Models based on X-ray crystal structures were constructed using PDB entries 4NPQ (GLIC, closed conformation) [6], 4HFI (GLIC, open conformation) [44], 3RQU (ELIC) [45], and 6FL9 (sTeLIC) [18]. Using Modeller [46] terminal residues to match the construct used in the SANS experiment were introduced to the models based on 4NPQ and 4HFI, as was residues missing in the 4NPQ structure. For the models based on 3RQU and 6FL9, homology models of GLIC with these structures as templates were constructed using Modeller. The N-terminal residues which were present in the construct but not in the crystal structures were visual inspected in the models, and a single conformation of these residues was chosen and introduced to all subunits in all models.

Models based on molecular dynamics (MD) simulations were obtained from simulations started from seeds along an open-closed transition pathway generated from coarse-grained elastic network-driven Brownian dynamics (eBDIMS) [30, 47]. Two transition pathways were constructed between X-ray structures 4NPQ and 4HFI resulting in a total of 50 seeds (25 for each pathway using either 4HFI or 4NPQ as eBDIMS target). The atomistic detail was then reconstructed using Modeller [46] in two steps; first, side chain conformations were transferred from 4HFI and energy minimized to fit the new conformation, followed by refinement of hydrogen bonds to ensure all secondary structure elements remained intact. Each initial conformation was then protonated to mimic both activating or resting conditions, resulting in a total of 100 initial conformations - 50 at activating and another 50 at resting conditions. For activating conditions a subset of acidic residues were protonated (E26, E35, E67, E75, E82, D86, D88, E177, E243; H277 doubly protonated) to approximate the predicted pattern at pH 4.6, as previously described [48]. Conformations were embedded in a 1-palmitoyl-2-oleoyl-sn-glycero-3-phosphocholine bilayer, soulibilized with TIP3P water and 0.1M NaCl, energy minimized, and equilibrated for 76 ns with gradual release of restraints. From the equilibrated systems, unrestrained MD simulations were launched, and frames after 200 ns of simulation time were extracted and used as models to fit against the SANS data. Terminal residues present in the experimental construct but not in the crystal structures were omitted in the simulations and extracted models. Energy minimization, equilibration, and MD simulations were performed using Gromacs [49] versions 2018.4 and 2019.3.

### 4.4 Data analysis

Guinier analysis was performed to calculate I(0) and radius of gyration (R_g_). In the Q-range from 0.010 Å^-1^ to 0.046 Å^-1^ (approximately 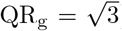) ln(I) was plotted as a function of Q^2^, to which a line was fitted. By applying the Guinier equation, the fitted line yielded I(0) from the intercept, and R_g_ from the slope. With I(0) determined, the molecular weight was calculated using

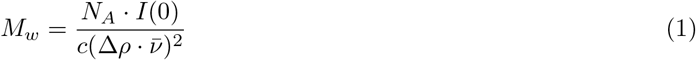

where N_A_ is Avogadro’s constant, c is the protein concentration, Δρ is the excess scattering length density, and 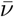 is the partial specific volume. The excess scattering length density is given by |ρ _protein_ - ρ_buffer_|, which was calculated using the scattering length density for GLIC and for D_2_O, see **Supplementary Table S3**. For GLIC the hydrogen-deuterium exchange of labile hydrogens was taken into account by calculating the expected degree of exchange after 100 minutes (approximately the time in the paused-flow experiments from the start of a run until the peak max) at pH 7.5 and pH 3.0 using PSX (Protein-Solvent Exchange) [50] and the model based on the closed GLIC structure. To account for GLIC being a transmembrane protein, which means that labile hydrogens in the transmembrane region are shielded to a larger extent than in a soluble protein, hydrogens within the transmembrane region as defined when submitting the model to the Positioning of Proteins in Membranes (PPM) webserver [51] were considered as ^1^H. The partial specific volume was estimated by dividing the volume of the closed pore model, calculated using ^3^V [52], by the molecular weight expected from the amino acid sequence.

The pair distance distribution was calculated from experimental curves using BayesApp [53] – available at https://somo.chem.utk.edu/bayesapp/ – and for the closed GLIC model using CaPP (Calculating Pair distance distribution functions for Proteins) [54]. Theoretical curves and fits to the experimental curves were calculated using Pepsi-SANS [55]. Linear combinations of theoretical curves (following the expression I_combination_ = k·I_model A_ + (1-k)·I_model B_ where I is intensity and k is the contribution from the first model) were tested using the curves calculated for the models of closed and open GLIC. The goodness of fit of the linear combinations to the experimental data was evaluated by calculating the reduced χ ^2^, approximating the degrees of freedom with the number of data points.

Principal component analysis (PCA) was performed using MDanalysis [56, 57], first constructing a PCA landscape using cartesian coordinates from the set of 46 GLIC structures used by Orellana et al. [30]. The models from MD simulations were projected onto this landscape and the correlation between χ^2^ and the first two principal components evaluated.

Graphs were plotted using Matplotlib [58], and protein images were rendered using Vmd [59]. A summary of software and equations used is available in **Supplementary Table S5**.

## 5 Data availability

The raw experimental data are available from:

Cuvette-SANS and continuous-flow SEC-SANS: https://doi.ill.fr/10.5291/ILL-DATA.8-03-959

Paused-flow SEC-SANS: https://doi.ill.fr/10.5291/ILL-DATA.8-03-1002

Processed SANS data and representative models are available at the following entries in the Small Angle Scattering Biological Data Bank (SASBDB): https://www.sasbdb.org/project/1317/

Cuvette-SANS at pH 7.5: SASDL63

Continuous-flow SEC-SANS at pH 7.5: SASDL53

Paused-flow SEC-SANS at pH 7.5: SASDL33

Paused-flow SEC-SANS at pH 3: SASDL43

## Acknowledgements

The authors acknowledge the Institut Laue-Langevin and ESRF for developments in SEC-SANS and allocated beam time at the D22 instrument. This work was supported by grants from SwedNess, the Knut and Alice Wallenberg Foundation, Swedish Research Council (2017-04641, 2018-06479, 2019-02433), Swedish e-Science Research Centre, BioExcel Center of Excellence (EU 823830), and from the Lundbeck Foundation through the Brainstruc grant. Computational resources were provided by the Swedish National Infrastructure for Computing (SNIC).

## Author contributions

Conceptualisation: ML, LA, RJH, EL; methodology: ML, CB, NTJ, RJH; software: ML; validation: ML; formal analysis: ML; investigation: ML, UR, CB, NTJ, AM, LP, RJH; resources: AM, LP, LA, RJH, EL; data curation: ML, CB, RJH, EL; original draft: ML, UR, CB, RJH; review & editing: ML, UR, CB, NTJ, AM, LP, LA, RJH, EL; visualization: ML, RJH; supervision: LA, RJH, EL; project administration: RJH; funding acquisition: EL.

