## Supplementary Information for "Probing solution structure of the pentameric ligand-gated ion channel GLIC by small-angle neutron scattering"

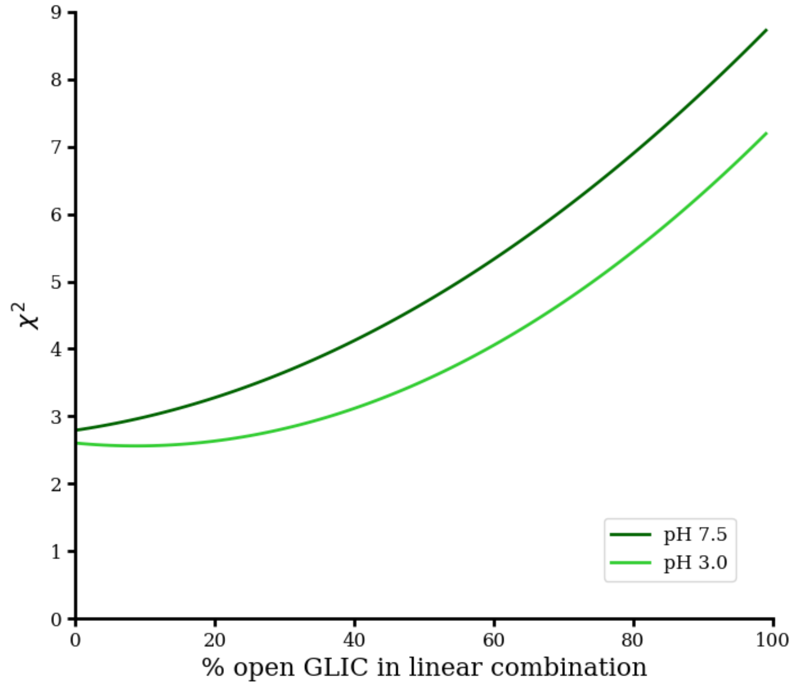


**Figure S1:** Goodness of fit ($\chi^2$) to SANS data collected under resting conditions (pH 7.5, dark green) or activating conditions (pH 3.0, light green), as a function of the contribution of the open GLIC crystal structure, in linear combination with the closed GLIC crystal structure.

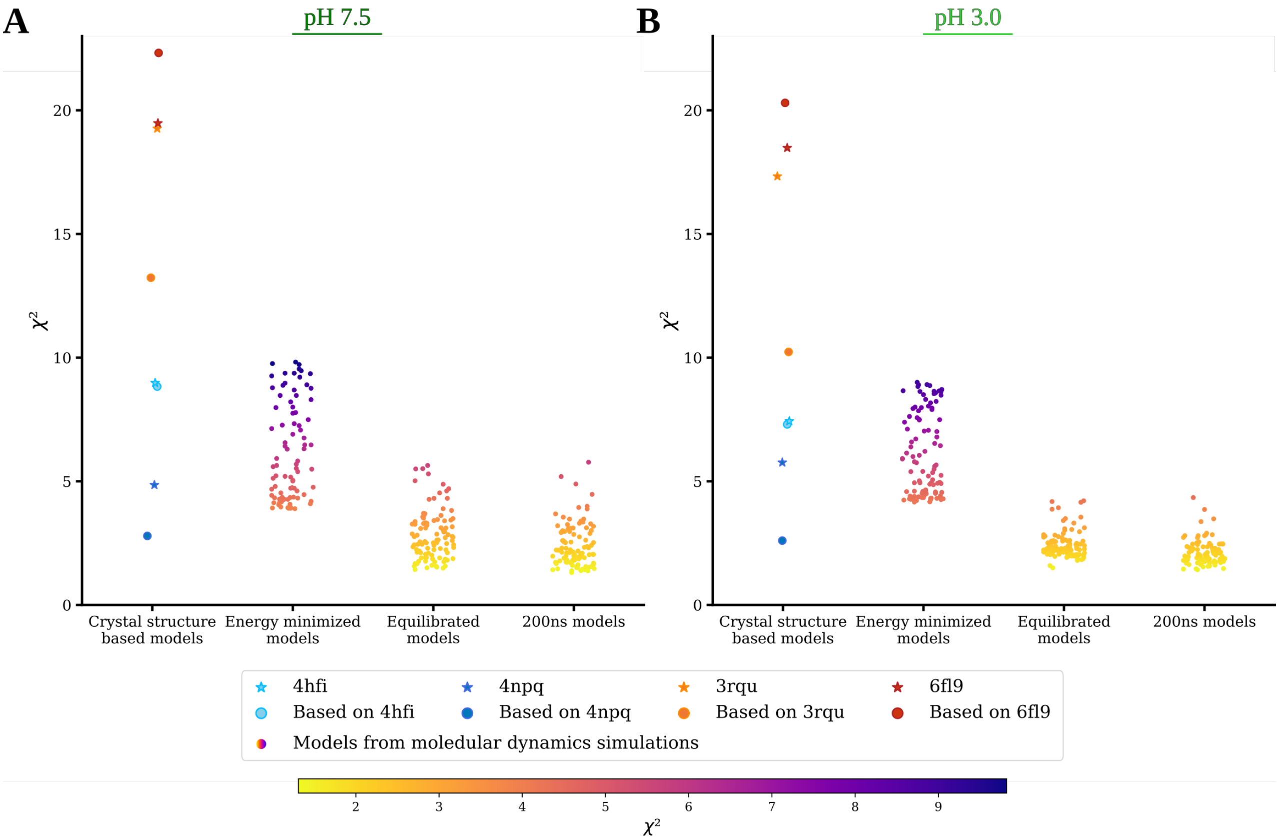


**Figure S2:** Comparison of model fits ($\chi^2$) to SANS data collected under **A)** resting (pH 7.5) or **B)** activating (pH 3.0) conditions. In the first column of each panel, crystal structures are shown as stars; models containing the full sequence of the processed GLIC construct are shown as circles, colored as for the crystal structures on which they were based. In subsequent columns, models from MD simulations are shown after energy minimization, after equilibration, and after 200 ns of unrestrained simulation, all as circles colored by $\chi^2$ value.

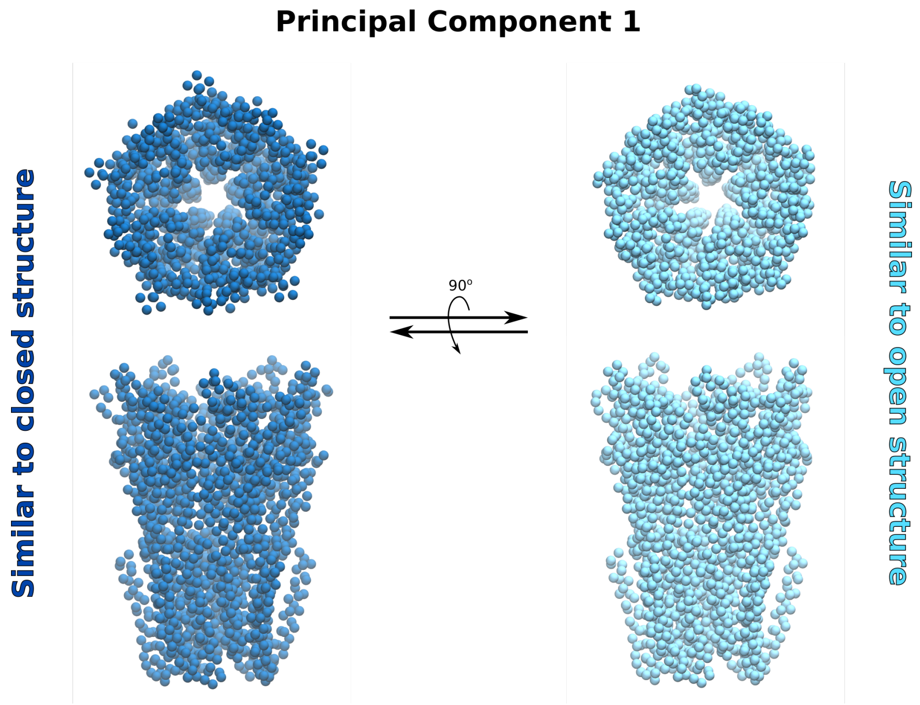


**Video S1:** The first principal component derived from alignment of 46 GLIC crystal structures as previously described [30], showing sequential projection of C$\alpha$ atom positions in energy-minimized models along the component. Models are shaded dark to light blue based on similarity to closed versus open crystal structures.

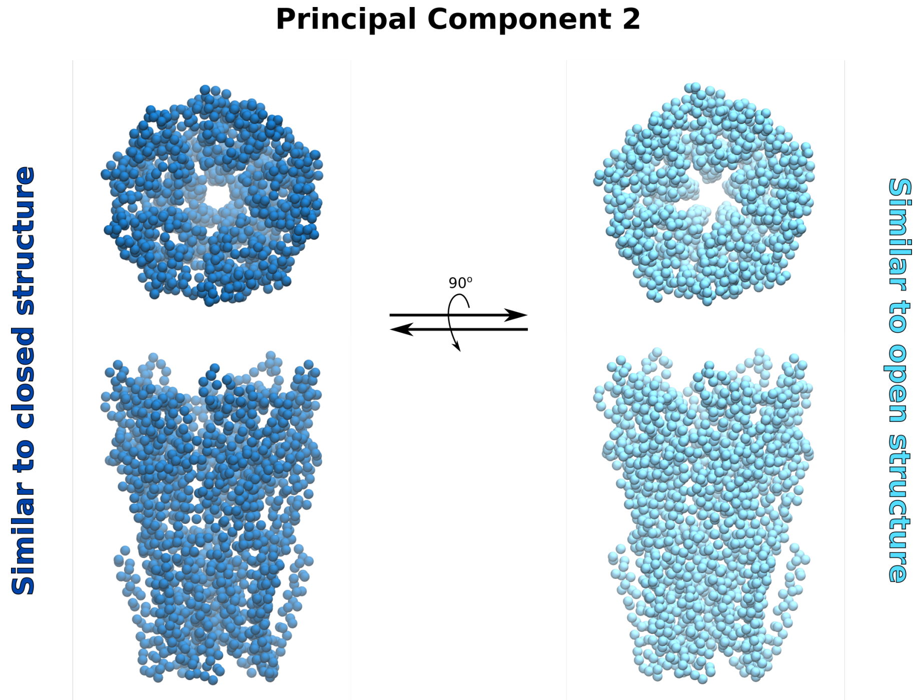


**Video S2:** The second principal component derived from alignment of 46 GLIC crystal structures as previously described [30], showing sequential projection of C$\alpha$ atom positions in energy-minimized models along the component. Models are shaded dark to light blue based on similarity to closed versus open crystal structures.

**Table S1:** Summary of structural parameters calculated from the paused-flow SEC-SANS data. For the Guinier analysis the minimum and maximum Q-value included is also listed.

| | pH 7.5 | pH 3 |
|---|---|---|
| Guinier analysis | | |
| $I(0)$ (cm$^{-1}$) | 0.44 | 0.45 |
| $R_g$ (Å) | 38.4 ± 0.2 | 38.3 ± 0.2 |
| $Q_{min}$ (Å$^{-1}$) | 0.010 | 0.010 |
| $Q_{max}$ (Å$^{-1}$), ($QR_g$ max) | 0.046, (1.7) | 0.046, (1.7) |
| Coefficient of correlation $R^2$ | 0.9994 | 0.9995 |
| $M$ from $I(0)$ (kDa), (ratio to predicted) | 194, (1.06) | 194, (1.06) |
| $P(r)$ analysis | | |
| $I(0)$ (cm$^{-1}$) | 0.441 ± .225E-03 | 0.445 ± .143E-03 |
| $R_g$ (Å) | 37.98 ± 0.03 | 37.81 ± 0.02 |
| $d_{max}$ (Å) | 109.50 ± 1.03 | 105.91 ± 0.54 |
| $q$ range (Å$^{-1}$) | 0.0060 - 0.2485 | 0.0060 - 0.2485 |
| $\chi^2$ | 1.04 | 0.91 |
| PepsiSANS | | |
| $R_g$ (Å) | 37.6 | 37.0 |

**Table S2:** Summary of modeling using protein structures and models based on protein structures, covering the radius of gyration of the model and the $\chi^2$ goodness of fit to the experimental data for the model.

| | | | | |
|---|---|---|---|---|
| Crystal structures | 4npq | 4hfi | 3rqu | 6fl9 |
| PepsiSANS | | | | |
| Predicted $R_g$ (Å) | 37.5 | 37.2 | 36.6 | 37.0 |
| pH 7.5 $\chi^2$ | 4.9 | 9.0 | 19.3 | 19.5 |
| pH 3.0 $\chi^2$ | 5.8 | 7.4 | 17.3 | 18.5 |
| All residue models | | | | |
| Based on | 4npq | 4hfi | 3rqu | 6fl9 |
| PepsiSANS | | | | |
| Predicted $R_g$ (Å) | 37.9 | 37.4 | 37.1 | 37.4 |
| pH 7.5 $\chi^2$ | 2.8 | 8.8 | 13.2 | 22.3 |
| pH 3.0 $\chi^2$ | 2.6 | 7.3 | 10.2 | 20.3 |
| MD-simulation models | | | | |
| eBDIMS | | | | |
| Extrapolation | From 4npq to 4hfi<br>From 4hfi to 4npq | | | |
| Output conformations (nr/extrapolation) | 25 | | | |
| Equilibration time (ns/simulation) | 76 | | | |
| Simulation time (ns/simulation) | 200 | | | |
| In aggregate (ns) | 20000 | | | |
| PepsiSANS | | | | |
| $\chi^2$ range | | | | |
| After energy minimization | | | | |
| pH 7.5 | 3.9 - 9.8 | | | |
| pH 3.0 | 4.2 - 9.0 | | | |
| After equilibration | | | | |
| pH 7.5 | 1.4 - 5.6 | | | |
| pH 3.0 | 1.5 - 4.2 | | | |
| After 200 ns simulation | | | | |
| pH 7.5 | 1.3 - 5.8 | | | |
| pH 3.0 | 1.4 - 4.3 | | | |

**Table S3:** Sample details, covering details of the protein, and experimental details like sample concentration, volume, and buffer.

| | |
|---|---|
| Organism of origin | *Gloeobacter violaceus* |
| Expression system | *Echerichia coli* |
| UniProt ID | Q7NDN8 |
| Extinction coefficient [$A_{280}$ 0.1%(w/v)] | 1.366 |
| Volume from structure (Å$^3$) | |
| All-residue model based on 4npq | 273150 |
| Particle contrast from sequence and solvent constituents, $\Delta\bar{\rho}$ ($\rho_{protein} - \rho_{solvent}$; $10^{10}$ cm$^{-2}$) | |
| pH 7.5 | 4.12 (2.25 - 6.37) |
| pH 3.0 | 4.14 (2.23 - 6.37) |
| M from chemical composition (kDa) | 182.7 |
| SEC-SANS | |
| Column | Superdex 200 Increase 10/300 |
| Loading concentration (mg/ml) | |
| Continuous-flow SEC-SANS | 4.0 |
| Paused-flow SEC-SANS | 5.6 |
| Injection volume (µl) | |
| Continuous-flow SEC-SANS | 300 |
| Paused-flow SEC-SANS | 240 |
| Flow rate (ml/min) | |
| Continuous-flow SEC-SANS | 0.3, 0.05 |
| Paused-flow SEC-SANS | 0.2, 0.01, 0 |
| Average concentration (mg/ml) in combined data frames | |
| Paused-flow SEC-SANS | |
| pH 7.5 | 0.77 |
| pH 3.0 | 0.74 |
| Cuvette SANS | |
| Concentration (mg/ml) | 0.47 |
| Volume (µl) | 360 |
| Solvent | |
| pH 7.5 | $D_2O$, 150 mM NaCl, 20 mM Tris·HCl, 0.5 mM d-DDM |
| pH 3.0 | $D_2O$, 150 mM NaCl, 20 mM citrate·HCl, 0.5 mM d-DDM |

**Table S4:** Summary the SANS collection parameters, including wavelength, pathlength, detector distances, Q-range, absolute scaling method, and normalization.

| | |
|---|---|
| Instrument | ILL D22 |
| Wavelength (Å) | 6 |
| Pathlength (cm) | |
| Cuvette SANS | 0.2 |
| Continuous-flow SEC-SANS | 0.1 |
| Paused-flow SEC-SANS | 0.1 |
| Detector distances (m) | |
| Cuvette SANS | 2m/2.8m & 11.2m/11.2m |
| Continuous-flow SEC-SANS | 2m/2.8m & 11.2m/11.2m |
| Paused-flow SEC-SANS | 2.8m/2.8m & 8m/8m |
| $Q$ measurement range ($Å^{-1}$) | |
| Cuvette SANS | 0.004 - 0.457 |
| Continuous-flow SEC-SANS | 0.004 - 0.456 |
| Paused-flow SEC-SANS | 0.006 - 0.452 |
| Absolute scaling method | Incident beam flux |
| Normalization | Divided by concentration |
| Exposure time (aggregate time) | |
| SEC-SANS | |
| Continuous-flow | 44 x 30s (22min) |
| Paused-flow, pH 7.5 | 116 x 30s (58min) |
| Paused-flow, pH 3.0 | 106 x 30s (53min) |
| Cuvette SANS | 2 x 1500s (50min) |
| Sample temperature (°C) | 10 |

**Table S5:** Summary of software and equations employed for SANS data reduction, analysis, and interpretation.

| | |
|---|---|
| SANS data reduction | GRASP v. 9.04 [42] |
| Extinction coefficient estimate | ProtParam [43] |
| Guinier equation | $\ln(I(Q)) = \ln(I(0)) - \frac{R_g^2}{3}Q^2$ |
| Calculation of $M_w$ | $M_w = \frac{N_A \cdot I(0)}{c(\Delta\rho \cdot \bar{\nu})^2}$ |
| Calculation of $\rho$ | $\rho = (\sum_{i=1}^{N} b_i)/V$ |
| Calculation of $\bar{\nu}$ | $\bar{\nu} = V/M_{aa}$ |
| Protein volume estimation | [3]V: Voss Volume Voxelator [52] |
| $P(r)$ analysis | BayesApp [53] *via* web server (https://somo.chem.utk.edu/bayesapp/) |
| $P(r)$ from structure | CaPP [54] (https://github.com/Niels-Bohr-Institute-XNS-StructBiophys/CaPP) |
| Atomic structure modelling | PepsiSANS v. 3.0 [55] |
| Missing sequence modelling | MODELLER v. 9.22 [46] |
| Structure extrapolation | eBDIMS [30, 47] |
| Molecular dynamics simulations | GROMACS v. 2018.4 and 2019.3 [49] |
| Theoretical $R_g$ | PepsiSANS v. 3.0 [55] |
| Hydrogen-deuterium exchange | PSX [50] |
| Three-dimensional graphic model representation | VMD [59] |
| Plots | MATPLOTLIB [58] |
